# Examining opioid withdrawal scoring and adaptation of global scoring systems to male and female C57BL/6J mice

**DOI:** 10.1101/2021.10.11.463944

**Authors:** Isabel Bravo, Maya Bluitt, Zoe McElligott

## Abstract

Opioid Use Disorder (OUD) is a chronic and relapsing psychiatric condition which is currently the leading cause of accidental death in the US. Symptoms of acute opioid withdrawal resemble a flu-like illness which is accompanied by a dysphoric state. Psychological comorbidities such as anxiety, depression, and disordered sleep can persist for months or years, well into the abstinence period. These symptoms are thought to drive further opioid intake in order to alleviate this unpleasant internal state. Many differences in OUD have been documented between male and female patients, with females at higher risk for relapse and overdose. This study sets out to characterize sex differences in symptoms and behavioral adaptations in mice during early withdrawal. Using our moderate dose, three-day precipitated withdrawal paradigm, we discovered significant effects of sex, time, and drug treatment on early withdrawal behaviors, locomotor activity, and gut motility in C57BL/6J mice. Here I will discuss previous methods of condensing behavioral phenotypes into one global withdrawal score, and propose a new methodology. This method increases the ability to detect nuanced effects and allows for more accurate translation across strain, sex, paradigm, and experimental context. Classification of opioid withdrawal-induced behavioral adaptations will allow for improved behavioral analysis of pharmacological manipulations, and investigations of brain circuitry involved in opioid withdrawal, as well as future screening of compounds with potential therapeutic benefit for the treatment of OUD.

---

Despite their known addictive potential, opioids such as morphine, oxycodone, and fentanyl are a widely prescribed group of analgesics. While opioids are highly efficacious for immediate pain relief, they are also associated with many adverse effects. The opioid epidemic in America has largely been driven by the over-prescription of legal opioids, which are often the first exposure for individuals who go on to develop OUD (CDC). Tolerance, or decreased efficacy of drug treatment over time, and opioid-induced hyperalgesia, or increased pain sensitivity, can drive patients to escalate drug use^1,2^.

Clinically, opioid withdrawal is accompanied by unpleasant physical and psychological symptoms which can drive disease relapse^3,4^. Our lab has developed a unique three-day paradigm as a clinically relevant animal model to study the effects of multiple bouts of opioid withdrawal in mice^5–7^. This paradigm makes use of a moderate dose of a general opioid receptor agonist and common clinical analgesic, morphine, over a shorter duration than other paradigms^8,9^. Naloxone, a competitive opioid receptor antagonist, used clinically to reverse opioid overdose, is administered to quickly precipitate withdrawal. We have previously demonstrated that this three-day paradigm produces early symptoms of withdrawal such as an increase in fecal boli production, escape jumping, paw tremors, and abnormal posture^7^. Additionally, we demonstrate significant behavioral adaptations in protracted withdrawal that have been observed 6-weeks after drug exposure, reminiscent of those reported clinically, which include changes in anxiety-like behavior, stress responsivity, sensitization to subsequent morphine challenge, and social interaction deficits^6^. Interestingly, we found sex differences in these features of protracted withdrawal^6^.

Much of the existing work investigating opioid withdrawal has been conducted exclusively in male rats. In order to study behavioral indications of altered internal state in our model of opioid withdrawal, we must verify that behaviors of interest are in fact indicative of changes due to morphine-naloxone pairing. In fact, this has been a challenge in clinical assessment of withdrawal, with many distinct scales for scoring opioid withdrawal in adults^10^ and infants^11^. The ability to then condense these behaviors into one global withdrawal score is of value to our research as it would allow for validation of treatment efficacy and examination of individual differences in response to morphine withdrawal. Here I evaluate previously developed global scoring systems and propose a new approach that would better express the state of withdrawal as a whole, and allow for identification of nuanced effects, including those related to treatment, time, and sex at lower sample sizes.

## Behavioral Adaptations in Early Precipitated Withdrawal

Validating and improving our model of opioid withdrawal is a continuous effort. Studying withdrawal in the context of sex differences provides new challenges, as classical withdrawal scales were developed using only male animals, specifically in male rats. Unfortunately, many differences in these withdrawal behaviors exist between these species and sexes that do not allow for easy translation. For example, we have not observed diarrhea or penile erection/ejaculation in our mice, especially not in females, so increased fecal boli production and genital grooming have been used as proxies, with the assumption that these behaviors are equivalent. These assumptions, however, have not been explicitly validated. Therefore, work is needed to probe for these sex differences and develop a scoring system that reflects early withdrawal accurately in both male and female mice.

To determine which behaviors accurately reflect a state of morphine withdrawal, mice were administered a subcutaneous injection of (0.9%) sterile saline (0.1 ml/10g) or 10 mg/kg morphine (in sterile saline 0.1 ml/10 g). All mice then received a subcutaneous injection of 1 mg/kg naloxone (in sterile saline 0.1 ml/10 g) to precipitate withdrawal two hours later. Freely behaving mice were observed in an open field environment during the 10 minutes following naloxone administration as previously described (Fig. 1)^6,7^. This paradigm has previously been shown to induce morphine dependence and subsequently withdrawal over the three-day administration period^7^.

**Figure 1.**
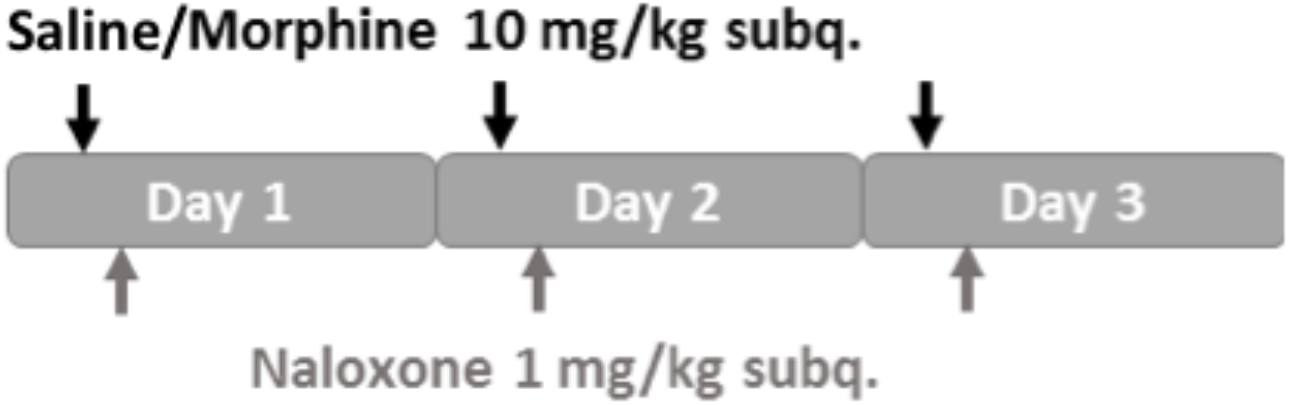
Morphine administration protocol. Mice were administered a subcutaneous injection of (0.9%) sterile saline (0.1 ml/10g) or 10 mg/kg morphine (in sterile saline 0.1 ml/10 g). All mice then received a subcutaneous injection of 1 mg/kg naloxone (in sterile saline 0.1 ml/10 g) to precipitate withdrawal two hours later, and were observed for withdrawal symptoms.

It was determined that escape jumps, paw tremors, abnormal posturing, and fecal boli production were significantly more pronounced in individuals of both sexes who had received morphine (Fig. 2a-c). Wet dog shakes and genital grooming were not found to be significantly increased during morphine withdrawal in our mice (Fig. 2d-e), and were therefore removed from subsequent scoring. In fact, genital grooming was decreased in morphine withdrawn mice, which we interpret as an anhedonic response and would expect to find an overall decrease in all types of grooming while mice experience this dysphoric internal state. Measures of gut motility were also taken, and it was determined that fecal boli number increased in morphine treated mice in comparison to their non-withdrawn counterparts (Fig. 2f-g). Furthermore, a time by drug treatment interaction was identified for weight wherein drug treated animals lost significant body mass over the three-day withdrawal period. Finally, locomotor behavior was also impacted by drug treatment, with withdrawn animals showing decreased time spent mobile and decreased average velocity during the 10-minute scoring period as compared to control animals (Fig. 2 h-i). In all, increases in escape jumping, paw tremors, abnormal posturing, and fecal boli production, as well as decreased locomotion can be used to measure the effects of drug treatment on early withdrawal symptoms. Identification of these behaviors and confirmation of their validity as measures of morphine withdrawal in this paradigm will be significant for future experiments examining the modulation of withdrawal by administration of various pharmacologic compounds.

**Figure 2.**
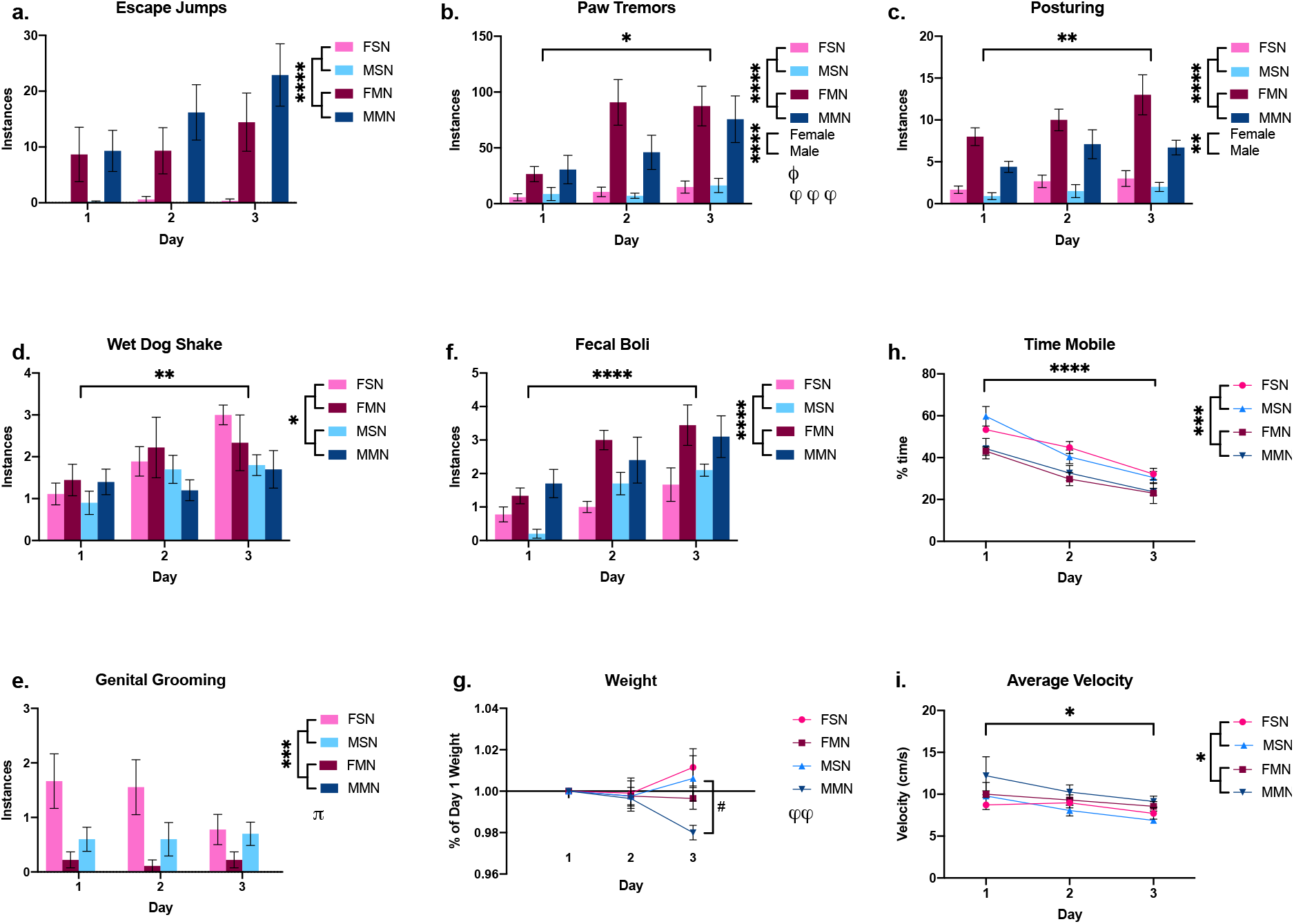
Morphine withdrawal alters behavior, locomotion, and gut motility. Groups: Female Saline-Naloxone (FSN), Female Morphine-Naloxone (FMN), Male Saline-Naloxone (MSN), Male Morphine-Naloxone (MMN). a. Withdrawal-induced escape jumping [time: F (1.915, 65.13) = 2.932, sex: F (1, 34) = 1.062, treatment: F (1, 34) = 28.98, time x sex: F (2, 68) = 0.4386, time x drug treatment: F (2, 68) = 2.878, sex x drug treatment: F (1, 34) = 1.262, time x sex x drug treatment: F (2, 68) = 0.6117.] Effect of drug treatment: ****p<.0001. b. Paw tremoring instances increase with morphine-naloxone pairing [time: F (1.943, 66.07) = 16.11, sex: F (1, 34) = 0.6919, drug treatment: F (1, 34) = 22.42, time x sex: F (2, 68) = 3.279, time x drug treatment: F (1, 34) = 0.7397, time x sex x drug treatment: F (2, 68) = 1.919.] Effect of drug treatment: ****p<.0001. Effect of time: ****p<.0001. Drug treatment x time interaction: ϕϕϕ p=0.0002. Sex x time interaction: Φ p=0.0437. c. Abnormal posturing increases in precipitated withdrawal [time: F (1.934, 65.76) = 6.974, sex: F (1, 34) = 10.35, drug treatment: F (1, 34) = 58.69, time x sex: F (2, 68) = 0.9133, time x drug treatment: F (2, 68) = 1.737, sex x drug treatment: F (1, 34) = 4.070, time x sex x drug treatment: F (2, 68) = 0.9546.] Effect of drug treatment: ****p <.0001. Effect of sex: **p=0.0028. Effect of time: **p=0.0020. d. Wet dog shake instance are unchanged but sexually dimorphic [time: F (1.866, 63.44) = 6.214, sex: F (1, 34) = 5.905, drug treatment: F (1, 34) = 0.005423, time x sex: F (2, 68) = 0.9899, time x drug treatment: F (2, 68) = 1.024, sex x drug treatment: F (1, 34) = 0.005423, time x sex x drug treatment: F (2, 68) = 0.8150.] Effect of sex: *p=0.0205. Effect of time: **p=0.0042. e. Genital grooming decreases with drug treatment [time: F (1.924, 65.40) = 1.635, sex: F (1, 34) = 4.082, drug treatment: F (1, 34) = 16.53, time x sex: F (2, 68) = 2.659, time x drug treatment: F (2, 68) = 2.049, sex x drug treatment: F (1, 34) = 1.381, time x sex x drug treatment: F (2, 68) = 3.220.] Effect of drug treatment: ***p=0.0003. Drug treatment x sex x time interaction: π p=.0461. f. Fecal boli production increases in morphine withdrawal [time: F (1.857, 63.14) = 16.24, sex: F (1, 34) = 0.0001995, drug treatment: F (1, 34) = 22.92, time x sex: F (2, 68) = 0.04952, time x drug treatment: F (2, 68) = 0.2498, sex x drug treatment: F (1, 34) = 0.5188, time x sex x drug treatment: F (2, 68) = 2.193.] Effect of drug treatment: ****p<.0001. Effect of time: ****p<.0001. g. Male morphine-treated mice lose weight in comparison to saline counterparts [time: F (1.776, 60.37) = 0.2372, sex: F (1, 34) = 1.053, drug treatment: F (1, 34) = 3.385, time x sex: F (2, 68) = 1.540, time x drug treatment: F (2, 68) = 5.770, sex x drug treatment: F (1, 34) = 0.2153, time x sex x drug treatment: F (2, 68) = 0.4541.] A priori t-test for sex differences (−0.02617 ± 0.01147), effect of drug treatment: # p=0.0350. 3-way ANOVA, treatment x time interaction: ϕϕ p=0.0048. h. Time spent mobile decreases in morphine treated mice [time: F (2, 30) = 56.16, sex: F (1, 15) = 0.1082, drug treatment: F (1, 15) = 19.07, time x sex: F (2, 30) = 0.7103, time x drug treatment: F (2, 30) = 0.6858, sex x drug treatment: F (1, 15) = 0.09068, time x sex x drug treatment: F (2, 30) = 1.073.] Effect of drug treatment: ***p=0.0006. Effect of time ****p<.0001. i. Average velocity decreases with morphine withdrawal [time: F (2, 30) = 4.947, sex: F (1, 15) = 0.6009, drug treatment: F (1, 15) = 5.646, time x sex: F (2, 30) = 1.069, time x drug treatment: F (2, 30) = 0.09265, sex x drug treatment: F (1, 15) = 1.262, time x sex x drug treatment: F (2, 30) = 0.03898.] Effect of drug treatment: *p=.0312. Effect of time: *p=.0139. For a-g: FSN and FMN n=9, MSN and MMN n=10. For h and I FSN n=4, and FMN, MSN and MMN n=5.

We have previously shown that early withdrawal related behaviors increase over the three days of our withdrawal paradigm^7^. As expected, many withdrawal-induced behaviors escalated over the three-day period, indicative of sensitization to the morphine-naloxone pairing. Paw tremors increased over time and exhibited a time x drug treatment interaction. Abnormal posturing, paw tremors, and fecal boli production also sensitized over the treatment period but did not show a time x drug treatment interaction. However, this increase is likely driven mainly by the morphine-treated group. Interestingly, wet dog shakes increased over time without an effect of drug treatment, however this difference looks to be driven mainly by females. Additionally, time spent mobile and average velocity both decreased over the three-day treatment period, potentially as a result of a reduction in exploratory behavior due to habituation to the novel environment. Since opioid use disorder (OUD) is a chronic and relapsing condition, it is important to validate that our animal model represents the cyclic and escalating nature of morphine dependence.

Sex differences in OUD have been well documented ^12–16^, and some work has also been done to examine sex differences in rodent models of OUD^14,17,18^. Our lab has previously found sex differences in inhibitory (GABA) synaptic transmission in bed nucleus of the stria terminalis (BNST) 24-hours following our three-day withdrawal paradigm as well as sex differences riod^6,7^. When we compared early withdrawal related behaviors in male and female mice, we found sex difference also exist at this timepoint. Paw tremors and posturing both showed an effect of sex, but not a sex x drug treatment interaction. It is possible that subtle differences exist in baseline behavior, or that naloxone treatment, regardless of if the animal received morphine, is responsible for increased instances in females. However, we hypothesize that this effect is driven by the morphine treated animals who show a more dramatic difference between males and females. Additionally, paw tremors show a significant sex x time interaction, as the females reach peak expression of the behavior earlier than their male counterparts. Further, genital grooming shows a significant sex x time x drug treatment interaction. This supports the hypothesis that grooming behavior may be an informative behavior, worthy of further investigation. Finally, because we had an a priori hypothesis that there would be sex differences in weight loss, we ran statistical analysis individually on males and females individually and found that male mice treated with morphine lost significantly more weight than their saline counterparts, while females showed no difference, consistent with our previous report^7,19^.

Due to a historical bias in the rodent literature, where many experiments have been performed exclusively in male subjects, it was important to develop a scoring system that would take sex differences in withdrawal presentation into account. Particularly, it needed to represent behaviors exhibited by both groups in order to be able to make fair comparisons between sexes so that we can determine if generalization of conclusions to both sexes is appropriate. In this case we found that while our paradigm induces significant behavioral changes immediately after precipitation of withdrawal, nuanced differences in behavioral expression exist between sexes.

## Re-examining global withdrawal scoring

There are several examples of scoring systems developed for use in male rats that allow individual behaviors to be collapsed into one global withdrawal score ^7^. These systems assigned a range for each behavior of interest with an associated point value, such that the total of the points would represent a global withdrawal score. However, in order to apply this method to a mouse model, it would have to be adapted in order to account for differences in types and frequency of behaviors between species. There is insufficient research on inter-species naloxone-precipitated morphine withdrawal differences to support this kind of manipulation, and would also require comparative assignment of values to distinct behaviors based on their significance to the overall withdrawal state, which would involve a great deal of speculation.

Our group has previously used a method for condensed global scoring wherein each escape jump is worth 1 point, and the presence of other withdrawal-related behaviors is awarded 1 point ^7^. Using the previously developed method of condensing withdrawal scores (Fig. 3a), it would appear as though both male and female animals who received morphine exhibit greater withdrawal associated behavior when compared to their saline associated counterparts. These behaviors also sensitize over the three sessions of exposure. This method of globalization assumes that escape jumps are the most reliable measure of withdrawal as they are given considerably more weight than any other behavior. Indeed, escape jumps are a well-documented feature of precipitated withdrawal in mice, however many other behaviors contribute to the overall expression of withdrawal state. Consistent with our previous study^7^, no effect of sex was observed on number of withdrawal jumps, however there were significant sex differences in other symptoms measured. Therefore, by minimizing the contribution of other withdrawal behaviors, this system would be unable to capture nuanced effects and interactions, especially those involving sex.

**Figure 3.**
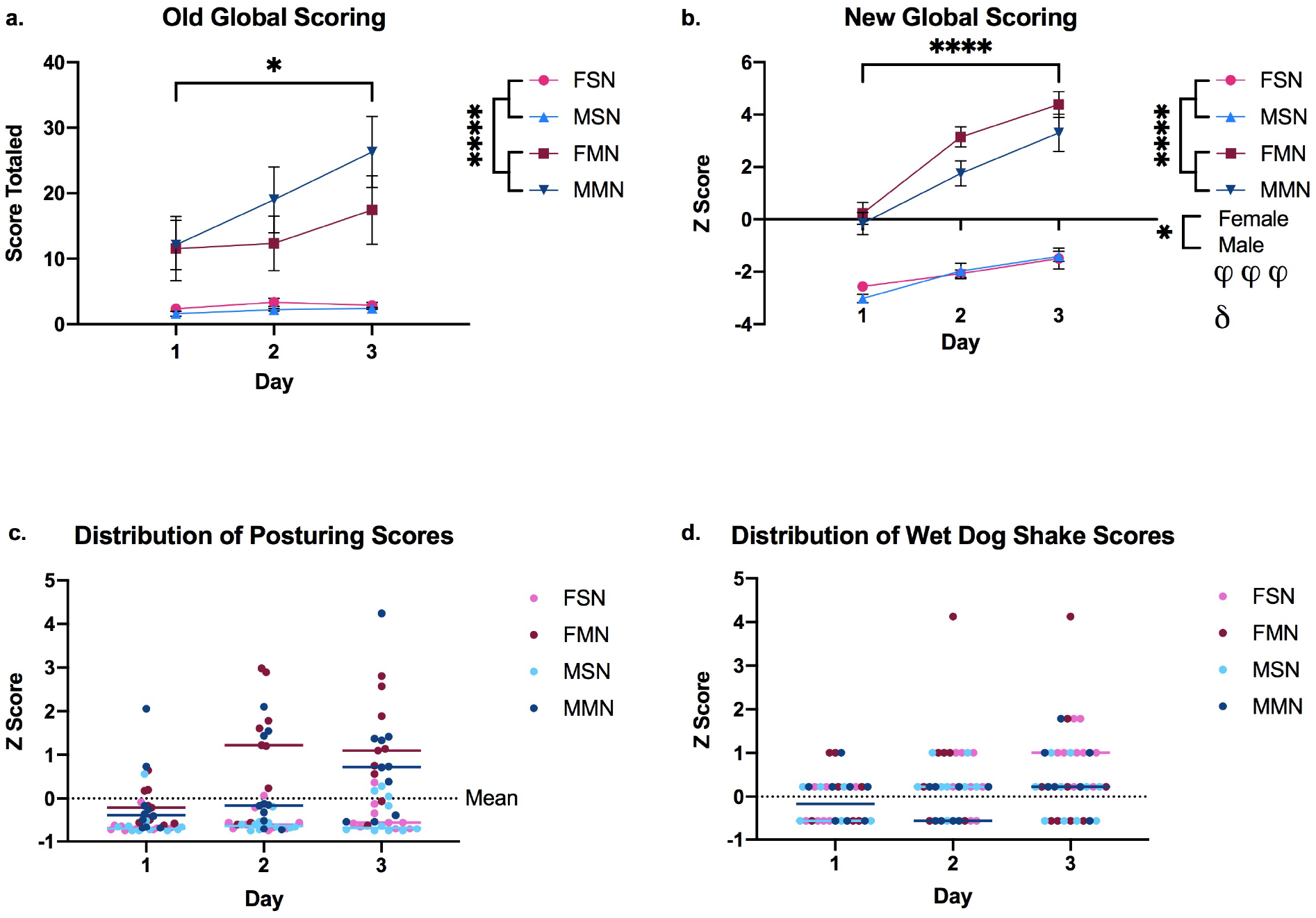
Understanding global withdrawal scoring using z-score system. a. Combined withdrawal scores with previously used model [time: F (2, 68) = 3.556, sex: F (1, 34) = 0.8537, drug treatment: F (1, 34) = 32.03, time x sex: F (2, 68) = 0.5840, time x drug treatment: F (2, 68) = 2.808, sex x drug treatment: F (1, 34) = 1.541, time x sex x drug treatment: F (2, 68) = 0.5637.] Effect of drug treatment: ****p<0.0001. Effect of time: *p=.0340. b. Combined withdrawal scores using z-score system [time: F (2, 68) = 40.77, sex: F (1, 34) = 6.148, drug treatment: F (1, 34) = 396.7, time x sex: F (2, 68) = 0.07988, time x drug treatment: F (2, 68) = 9.539, sex x drug treatment: F (1, 34) = 4.187, time x sex x drug treatment: F (2, 68) = 1.023.] Effect of drug treatment: ****p<0.0001. Effect of sex: *p=.0183. Effect of time: ****p<0.0001. Drug treatment x time interaction ϕϕϕ p=.0002. Drug treatment x sex interaction: δ p=.0485. c. Example of distribution of z-scores for a behavior (paw tremors) with effect of drug treatment.

As a solution to this escape jump biasing in global withdrawal score, I propose that each of the behaviors determined to be indicative of morphine withdrawal be given weight based on the strength of the difference expressed (Fig. 3b). That is to say that if two groups are treated, one with morphine and the other with vehicle, then behaviors that are more modulated by drug treatment will exhibit greater variation between treatment groups (Fig. 3c) than ones with only a slight effect or no effect (Fig. 3d). In order to account for the differences in score range between behaviors, each subject’s score is represented as a standardized measurement of variance, in this case a z-score, or the number of standard deviations an individual animal’s score falls from the average of observed scores for that behavior across all groups and days of treatment. Individuals within a treatment group would naturally cluster together above or below the average if the features of that group (sex and/or drug treatment) affected expression of that behavior. Additionally, the effect of time on behavioral score could also be accounted for with this method. If a behavior sensitizes over the 3-day period, we would expect that on day 1 scores would be below average, or receive a negative z-score, while on day 3 they would be above average, represented by a positive z-score. Examining two examples, these trends can be visually understood. A behavior, like wet dog shakes, that shows no significant effect of drug treatment produces a homogenized plot where morphine and saline groups of respective sexes occupy the same space (Fig. 3d). In contrast, paw tremors show a robust effect of drug treatment, time, time x drug treatment interaction, and sex x time interaction. As these trends develop, treatment groups become more distinct and the plot becomes heterogeneous, representing these differences (Fig. 3c). When taken as a whole, weighted based on contribution to the altered internal state produced by our paradigm of opioid withdrawal, we can see that drug treatment, sex, and number of bouts of exposure all have interesting and intertwined relationships that are expressed through the behaviors we have identified here.

## Conclusions

Here we demonstrate that there are many behavioral, locomotor, and gut-related effects of our paradigm of morphine withdrawal. In order to better understand one aspect of withdrawal, early behavior phenotypes, scores for individual behaviors can be condensed into a global withdrawal score. By representing behavior scores as the number of standard deviations from the mean of that behavior, each behavior is weighted by the extent to which expression is changed by an independent variable. This method can be easily adapted to different withdrawal manipulations, as the behavior of control mice is used as a comparison, as opposed to assuming that mice will perform the same behaviors regardless of context. Providing set cutoffs for the assignment of point values, as has been done previously, only works within the context and conditions in which these values were originally assigned. Strain, species, sex, age, experimenter, environment, dose, rout of administration, time of testing, and cage environment can all change behavioral output. As an example, addition of a bedding to a box can encourage digging and reduce time for other behaviors being scored. Anecdotally, our lab has observed that administering morphine and naloxone in the same context prevents escape jumping, whereas administration of morphine in the home environment and naloxone in a novel one produces vigorous jumping. Mouse strains with high baseline anxiety or locomotion exhibit vastly different behavioral phenotypes than their slower counterparts as I have observed working with strains of mice in the Collaborative Cross project.

It is unlikely that when created, early withdrawal scoring systems were meant to be used as the definitive scale for withdrawal behavior assessment, but rather were adopted in the absence of more appropriate tools. Rather than assume that findings that do not reproduce original findings are incorrect, this new, flexile method of using z-scores allows for the identification of unanticipated findings and elaboration on current knowledge. This new scoring system will be used in the future to understand how individual variation in behavior corresponds with changes in brain activity, to compare the effects of norepinephrine modulating drugs on opioid withdrawal, and to screen compounds like oxytocin for therapeutic potential. One limitation to consider is that this global scoring system has only been tested in one model of opioid withdrawal. Subsequent experiments will be performed to test this system in a paradigm of fentanyl drinking to establish translational value. Further work should be done to make sense of the interesting changes in grooming behavior observed as well as expand on our understanding of gut motility and locomotor tendencies as they relate to the dysphoria associated with opioid withdrawal.

## Methods

All procedures were approved by the University of North Carolina Institutional Animal Use and Care Committee.

### Subjects

Male and female, 8 to 12-week old C57/BL6 mice obtained from Jackson Laboratories were group or signally housed and habituated to their home-cage environment for 2 weeks prior to all manipulations. Mice were housed on a 12:12 hour light-dark cycle and provided access to food and water ad libitum. Prior to behavioral experiments, mice received two sessions of handling to minimize later stress, and allowed to recover undisturbed for one week. A subset of mice scored for withdrawal symptoms (20) were exposed to Ensure in their home cage for 3 hours on 4 consecutive days, one week prior to withdrawal for a later experiment. The remaining were exposed to the scent of vanilla during withdrawal. These manipulations were necessary for subsequent experiments performed on these cohorts, but should not significantly impact data shown here.

### Morphine Withdrawal

On withdrawal days, as previously described in Luster, et al. (2019) and Bravo, et al. (2020), mice received a subcutaneous injection of (0.9%) sterile saline (0.1 ml/10g) or 10 mg/kg morphine (in sterile saline 0.1 ml/10 g). All mice then received a subcutaneous injection 1 mg/kg naloxone (in sterile saline 0.1 ml/10 g) in a distinct environment two hours later, and were observed for withdrawal symptoms over the following 10 minutes. A subset of these mice (20) were exposed to the scent of vanilla (in glycerol) contained in an adapted conical tube in the corner of the observation arena as described above. All mice where observed for symptoms 10 minutes following withdrawal, and videos were taken of the scent receiving animals, which was later assessed for locomotor activity using BehaviorCloud software.

### Statistics

Withdrawal data were analyzed using 3-way ANOVA. Statistical significance was set at α = 0.05, and exact values are provided in figure legends. A priori t-tests for sex differences were conducted when appropriate. The statistical analyses for behavioral experiments were performed using GraphPad Prism 9 software. Two mice were excluded from the study, one received a small needle stick in the foot during injections and the other disappeared from its cage in the animal facility.

